# Restoration of visual function by transplantation of optogenetically engineered photoreceptors

**DOI:** 10.1101/399725

**Authors:** Marcela Garita-Hernandez, Maruša Lampič, Antoine Chaffiol, Laure Guibbal, Fiona Routet, Tiago Santos-Ferreira, Giuliana Gagliardi, Sacha Reichman, Serge Picaud, José-Alain Sahel, Olivier Goureau, Marius Ader, Deniz Dalkara, Jens Duebel

**Affiliations:** Sorbonne Université, Institut de la Vision, INSERM, CNRS, 75012 Paris, France; CRTD/Center for Regenerative Therapies Dresden, CMCB, TU Dresden, Germany; CHNO des Quinze-Vingts, DHU Sight Restore, INSERM-DGOS CIC 1423, Paris, France; Department of Ophthalmology, The University of Pittsburgh School of Medicine, USA

## Abstract

A major challenge in the treatment of retinal degenerative diseases, with the transplantation of replacement photoreceptors, is the difficulty in inducing the grafted cells to grow and maintain light sensitive outer segments (OS) in the host retina, which depends on proper interaction with the underlying retinal pigment epithelium (RPE). For a RPE-independent treatment approach, we introduced a hyperpolarizing microbial opsin into photoreceptor precursors from new-born mice, and transplanted them into blind mice lacking the photoreceptor layer. These optogenetically transformed photoreceptors were light responsive and their transplantation lead to the recovery of visual function, as shown by ganglion cell recordings and behavioral tests. Subsequently, we generated cone photoreceptors from human induced pluripotent stem cells (hiPSCs), expressing the chloride pump Jaws. After transplantation into blind mice, we observed light-driven responses at the photoreceptor and ganglion cell level. These results demonstrate that *structural* and *functional* retinal repair is possible by combining *stem cell therapy* and *optogenetics*.

---

Cell replacement therapy offers hope for the treatment of late stage retinal degeneration, when the outer retinal photoreceptor layer is lost ^1-3^. However, a remaining obstacle of photoreceptor replacement is that transplanted cells have to develop into functional photoreceptors with light sensitive OS. Indeed, in mouse models of severe degeneration, the formation of light-sensitive OS by transplanted photoreceptors has been difficult to achieve ^4-6^. Recent studies, using retinal sheet transplantation lead to major improvements in terms of OS formation and light sensitivity ^7, 8^. Despite these promising results, a major problem has not yet been solved: photoreceptors need tight interaction with the RPE in order to maintain their structure and function via continuous disc shedding and renewal ^9^. Since in retinal degenerative diseases the RPE is usually also compromised ^10^, the probability that transplanted photoreceptors stay sensitive to light is very low ^11, 12^. To tackle this problem, we introduced optogenetic light sensors into photoreceptors, derived from the developmental mouse retina as well as from hiPSCs, and transplanted them into mouse models of severe retinal degeneration. The key point of our approach is that these optogenetically transformed photoreceptors stay functional based on the activity of the microbial opsin, even in the absence of properly formed OS and without the support from the RPE.

For optogenetic transformation of mouse photoreceptors, eyes of new-born wild-type mice at post-natal day (P) 2 were injected with an adeno-associated viral (AAV) vector encoding enhanced *Natronomonas pharaonis* halorhodopsin eNpHR2.0 (NpHR) ^13^ under the control of the rhodopsin promoter (AAV-Rho-NpHR-YFP) (Fig. 1A and Fig. S1). At P4, photoreceptor precursors were magnetic activated cell (MAC) sorted by using the photoreceptor specific cell surface marker CD73 ^14, 15^. The harvested cells were transplanted via sub-retinal injections into two blind mouse models of late-stage retinal degeneration (*Cpfl1/Rho*^*-/-*^ ^16^, age 5 to 18 weeks; *C3H rd/rd* (*rd1)* ^17^, age 4 to 11 weeks). At this age, almost all of the outer nuclear layer (ONL) was lost in untreated mice (Fig. 1, B and E). Four weeks after transplantation, we investigated the morphology of the transplanted donor cells and their ability to integrate into the host retina. In both mouse models, we found NpHR-positive donor cells in close contact to cell bodies of rod bipolar cells, but none of the transplanted cells displayed correctly formed OS (Fig. 1, C,D,F,G). We tested if we can elicit light responses from NpHR-positive donor cells in the absence of functional OS. Two-photon targeted patch-clamp recording revealed robust responses to light pulses (580 nm, 10^16^ photons cm^-2^s^-1^) (Fig. 1H and Fig. S2). There were no measurable light-evoked currents in transplanted photoreceptors expressing GFP only, which is consistent with the finding that the transplanted cells lacked their light sensitive OS. Stimulation at different wavelengths showed a spectral sensitivity matching the action spectrum of NpHR (Fig. 1I). To measure the temporal properties of NpHR-positive photoreceptors, we recorded photocurrents using light pulses at increasing frequencies, and we observed that they could follow up to 25 Hz (Fig. 1J and Fig. S2). The rise constants were significantly faster compared to photocurrents of wild-type mice (Fig. 1K). Both, from the spectral (peak current at 580 nm) and the temporal (Tau _ON_ < 10ms) response properties we conclude that the photocurrents were driven by the introduced NpHR (Fig. 1, I-K, and fig. S2).

**Fig. 1.**
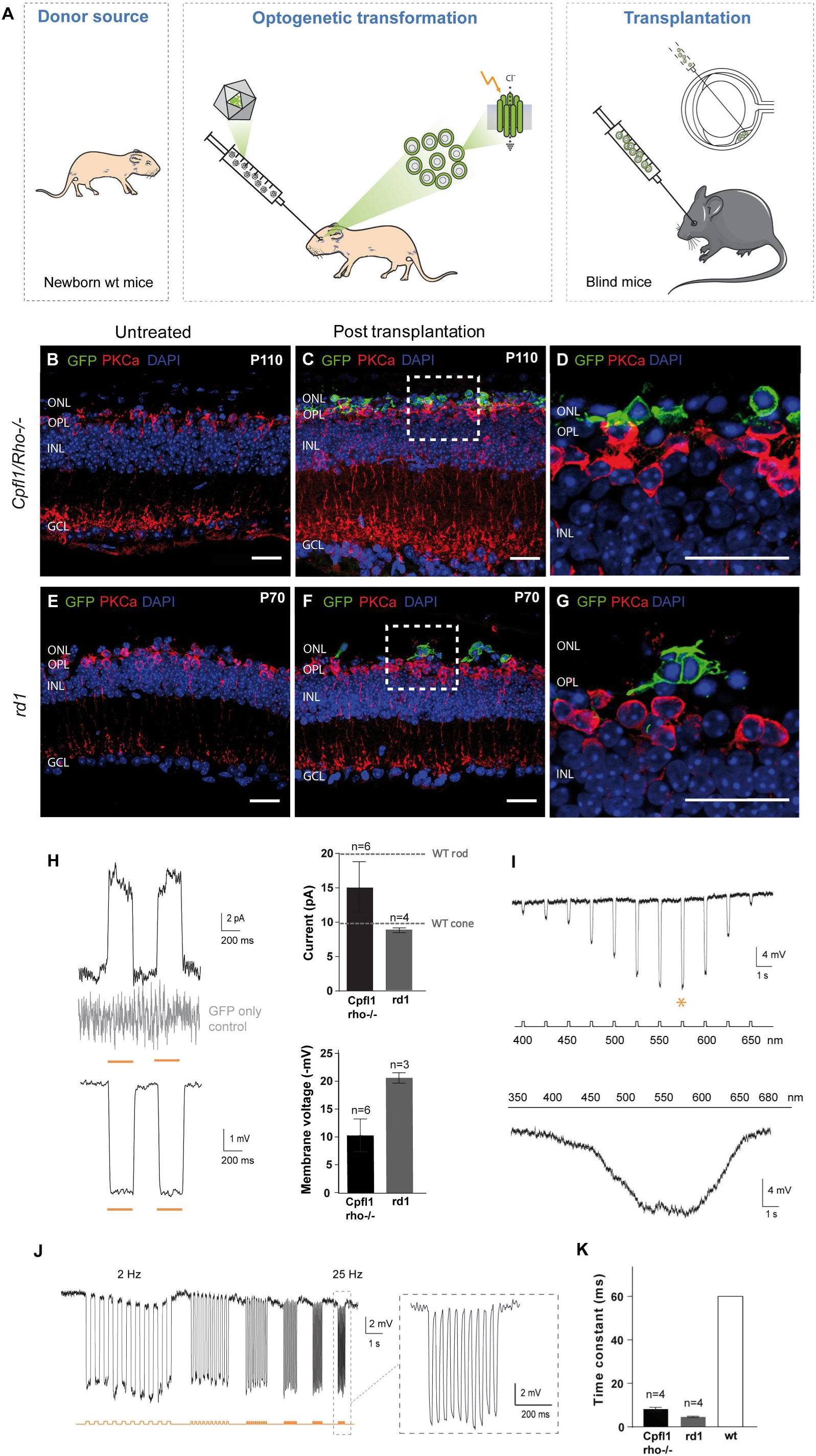
Transplanted photoreceptor precursors, expressing NpHR, integrate into the retina of blind mice and respond to light. **(A)** Eyes of wt mice at P2 were injected with AAV-Rho-NpHR-YFP. Two days later, retinas were dissected and photoreceptor precursors sorted out. These cells were transplanted via sub-retinal injections into blind mice. (B-G) Immunofluorescence analysis on vertical sections of *Cpfl1/Rho*^*-/-*^ (B-D) and *rd1* (E-G) retinas. **(B)** Age-matched non-transplanted *Cpfl1/Rho*^*-/-*^ retina. **(C,D)** *Cpfl1/Rho*^*-/-*^ retina transplanted with NpHR-photoreceptors showing YFP+ cells (green) located on top of host PKCα bipolar cells (red). **(E)** Age-matched non- transplanted *rd1* retina. **(F,G)** *rd1* retina transplanted with NpHR- photoreceptors. Scale bars 25 μm. (H-L) Light response characteristics from cells recorded by whole-cell patch-clamp technique in treated *Cpfl1/Rho*^*-/-*^ mice. **(H)** Left, light-evoked responses of NpHR- photoreceptors stimulated with 2 consecutive flashes (top, current response; bottom, voltage response), absence of the response in GFP-only expressing PR shown in grey. Right, comparison of response amplitudes. Mean photocurrent peak (top) and mean peak voltage response (bottom). Mean values observed in wt rods and cones are indicated with a dashed line ^25^. **(I)** Representative action spectrum from a NpHR- photoreceptors stimulated at different wavelengths. Top, stimuli ranging from 400 nm to 650 nm, separated by 25 nm steps. Maximal voltage responses were obtained at 575 nm. Bottom, continuous ‘rainbow’ stimulation between 350 and 680 nm. **(J)** Temporal properties: Modulation of NpHR-induced voltage responses at increasing stimulation frequencies from 2 to 25 Hz. (**K**) Comparison of rise time constants in the two models and in wild type cones. All light stimulations were performed at 1.3 10^16^ photons cm^-2^ s^-1^ and 590 nm, if not stated otherwise. In all panels: n means number of cells.

Next, we investigated if the signals from transplanted photoreceptors are transmitted to retinal ganglion cells, the output neurons of the retina. By using extracellular spike recording, we measured ON- and OFF-light responses in retinal ganglion cells. These results demonstrate that the transplanted photoreceptors can form synaptic connections with the inner retinal neurons and that NpHR-induced signals are transmitted to the retinal output neurons via ON- and OFF-pathways (Fig. 2A and Fig. S3). Recordings performed under pharmacological blockade of photoreceptor input to ON-bipolar cells (50 µM L-AP4) showed complete abolition of ON light responses, which recovered after 20 minutes of L-AP4- washout. These control experiments confirmed that light induced signals were indeed transmitted via photoreceptor-to-bipolar cell synapses (Fig. 2, B and C). By stimulating treated retinas at different wavelengths we determined the spectral sensitivity, and we observed peak responses at 550-600nm, reflecting the action spectrum from NpHR (Fig. 2, D and E). To assess the light intensities required to trigger spike responses, we used light pulses (580 nm) at different intensities. Importantly, the intensities required to evoke light responses were well below the safety limit for optical radiation in the human eye ^18, 19^ (Fig. 2, F and G, and Fig. S3). We did not observe measurable light responses in retinae from age-matched control-mice, where photoreceptors, expressing GFP only, were transplanted (Fig. 2, H and I, and Fig. S3). To test whether the behaviour of treated mice could be modulated by light, we used the light/dark box test ^20^ (Fig. 2J). Treated *Cpfl1/Rho*^*-/-*^ mice displayed a robust light avoidance behaviour (35.8±1.8% of time in the illuminated compartment), compared to non- injected (57.3±2.6%) and GFP-only expressing control mice (56.4±6.4%) (Fig. 2K).

**Fig. 2.**
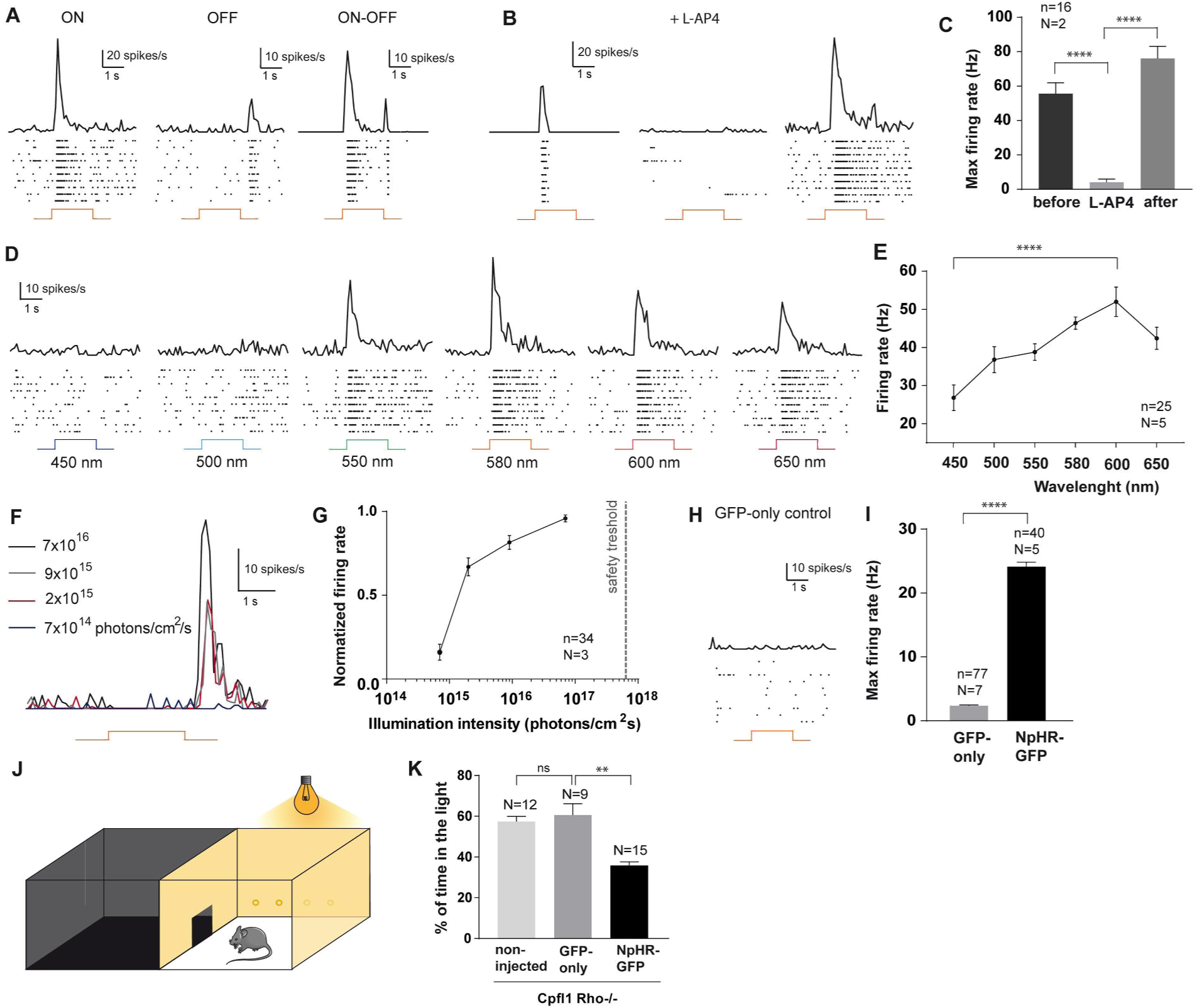
NpHR-triggered responses from transplanted photoreceptors are transmitted to retinal ganglion cells (RGC) and induce light avoidance behaviour in blind mice. (A-I) Averaged spike responses obtained from multi-electrode array (MEA) recordings shown as PSTH and raster plots recorded in transplanted *Cpfl1/Rho*^*-/-*^ mice (stimulation: 580 nm, 7× 10^16^ photons cm^-2^ s^-1^). **(A)** Representative traces from three RGCs responding either with an ON-, OFF-, or ON/OFF-response pattern. **(B)** Representative traces from a cell before, during ON bipolar cell blockade, and after wash-out, and **(C)** quantification of maximum firing rates for these conditions. **(D)** Representative responses to wavelengths ranging from 450 nm to 650 nm. **(E)** Quantification of RGC action spectrum. The cells reach their peak firing rate at 600 nm. **(F)** PSTHs of a single RGC responding to stimuli of increasing intensities (from 7 × 10^14^ to 7× 10^16^ photons cm^-2^ s^-1^). **(G)** Intensity curve. The dashed line indicates the maximum light intensitiy allowed in the human eye at 590nm ^18, 19^. **(H)** Unresponsive cell from a control retina transplanted with GFP-only-expressing photoreceptors. **(I)** Maximum firing rate in mice treated with GFP-only photoreceptors versus mice treated with NpHR- photoreceptors. **(J)** Schematic representation of the dark/light box test. **(K)** Percentage of time spent in the light compartment for: non-treated *Cpfl1/Rho*^*-/-*^ mice, *Cpfl1/Rho*^*-/-*^ mice treated with GFP-only photoreceptors, and *Cpfl1/Rho*^*-/-*^ mice treated with NpHR-photoreceptors (illumination: 590 nm, 2.11 × 10^15^ photons cm^-2^ s^-1^). In all panels: n, number of cells, N, number of mice; error bars, SEM; stars, statistical significance, ns, nonsignificant ^26^.

To evaluate the translatability to human subjects, we asked if it is possible to replace the mouse donor source with optogenetically-engineered hiPSCs (Fig. 3A). Therefore, we optimized a previous protocol of differentiation based on the self-generation of 3D neural- retina-like structures ^21^. Using this system, we generated cone-enriched retinal organoids, expressing the pan-photoreceptor markers CRX and recoverin (RCVRN) as well as the cone-specific marker cone arrestin (CAR) (Fig. 3, B-F and Fig. S4). Cone photoreceptors are responsible for high acuity daylight vision in humans, and are therefore the preferred choice for transplantation. To render these cones light sensitive, we used the hyperpolarizing chloride pump Jaws, based on its enhanced expression level and improved membrane trafficking in human tissue, compared to NpHR ^22^,^23^. By using an AAV vector, encoding Jaws-GFP under the control of CAR promoter, we delivered the microbial opsin to the hiPSC-derived cone photoreceptors (Fig. 3, G and H). Single cell recording of GFP-positive Jaws-expressing cones in retinal organoids revealed solid light responses, matching the response properties of Jaws, while recordings from hiPSC-derived cones, expressing GFP only, showed no light responses (Fig. 3, I-L). Additionally, monolayer cultures of human cones expressing Jaws, show no outer segments and strong light responses (Fig. S5). These results demonstrate that it is possible to induce robust optogenetic responses in photoreceptors derived from hiPSCs in the absence of light sensitive OS.

**Fig. 3.**
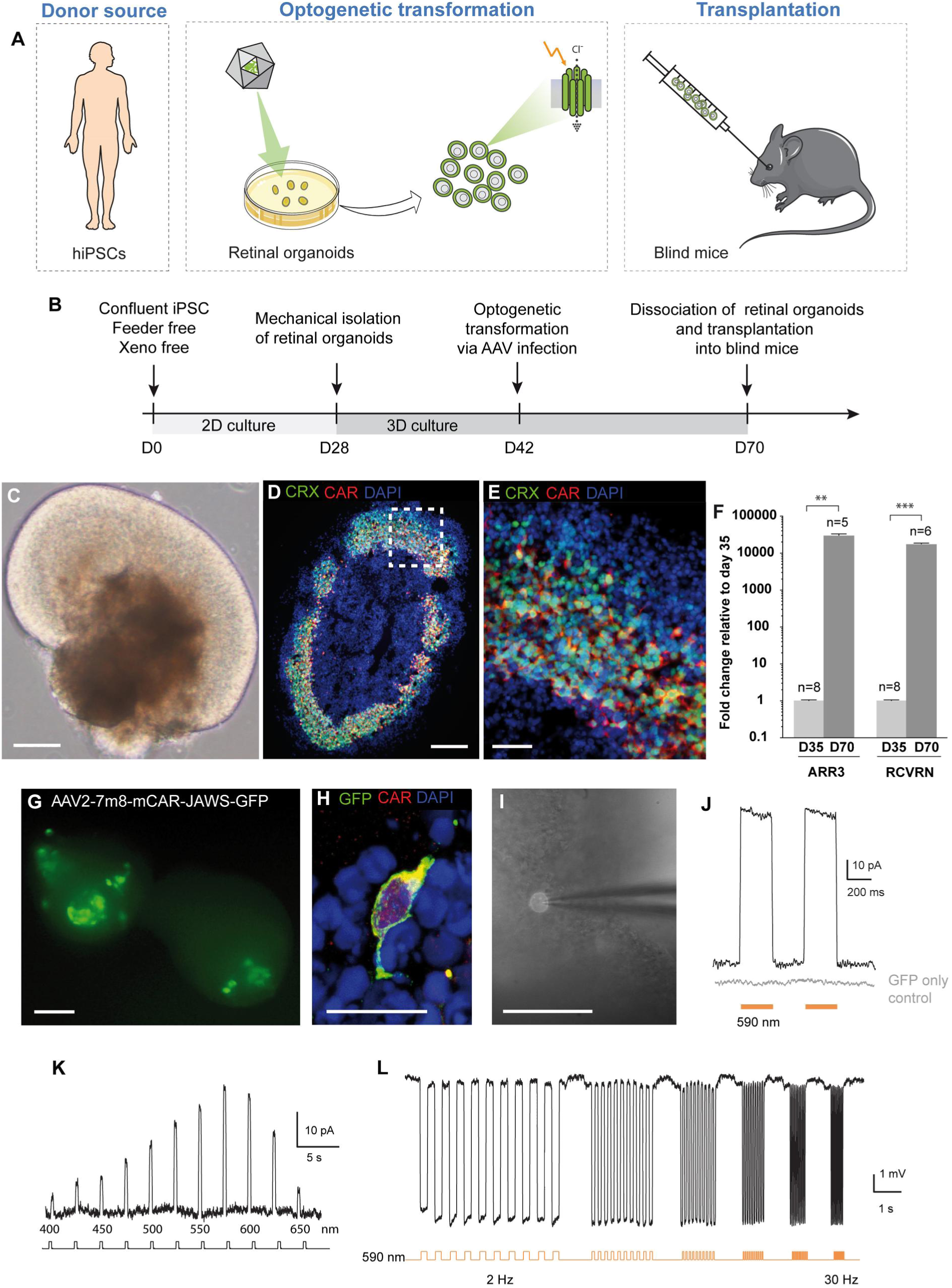
Jaws-expressing photoreceptors, derived from hiPSCs, are sensitive to light. **(A)** Human iPSCs were differentia ted towards retinal organoids and were infected with AAV-mCar-Jaws-GFP. After further maturation, cells were dissociated and iPSC-derived photoreceptors were transplanted into blind mice. **(B)** Schematic diagram of the differentiation and viral transformation of retinal organoids. **(C)** Bright-field image of a retinal organoid at D30 of differentiation. **(D, E)** Characterization of a representative retinal organoid at D70, depicting a thick layer of photoreceptors immunoreactive for CRX (green) and CAR (red). **(F)** Real time qRT-PCR analysis of photoreceptor specific markers *CAR (ARR3)* and *RCVRN*. n, number of biological replicates; error bars, SEM; stars, statistical significance ^26^. **(G)** Live GFP fluorescence observed at D54 (12 days post infection). **(H)** A single cone photoreceptor stained with GFP (green) and CAR (red) at D70. **(I)** Bright field/epifluorescence image of a GFP+ cell patched inside a retinal organoid at D70 of differentiation. Scale bars, C,D,G,I 100 μm; E,H 25 μm. (J-L) Patch-clamp data from Jaws- cones within organoids, stimulation at 590 nm if not stated otherwise. **(J)** Photocurrent responses after stimulation with 2 consecutive flashes at 3.5 10^17^ photons cm^-2^ s^-1^, absence of response in GFP-only expressing cones is shown in grey. **(K)** Photocurrent action spectrum corresponding to a Jaws-cone stimulated at wavelengths ranging from 400 nm to 650 nm. Maximal responses were obtained at 575 nm (at 1.3 10^16^ photons cm^-2^ s^-1^). **(L)** Modulation of Jaws-induced voltage responses at increasing stimulation frequencies from 2 to 30 Hz.

In order to transplant Jaws-positive photoreceptors, we dissociated the retinal organoids and injected the cell suspension subretinally into the blind hosts (*Cpfl1/Rho*^*-/-*^, age 10 to 15 weeks; *rd1*, age 4 to 5 weeks) (Fig. 3A). In both *Cpfl1/Rho*^*-/-*^ and *rd1* mice we observed Jaws- expressing donor cells in close proximity to the host INL (Fig. 4, A-D). The transplanted GFP-positive cells were located next to PKCα-positive bipolar cells (Fig. 4A) and the synaptic marker Synaptophysin was localized in close apposition to the bipolar cell dendrites (Fig. 4B), indicating that the human cells are able to form synaptic connections with the host bipolar cells. By using a human-specific nuclear marker (Human Nuclear Antigen, HNA) we confirmed the human origin of these cells (Fig. 4C), and the photoreceptor specific marker (RCVRN) confirmed that the transplanted cells, derived from hiPSCs, were differentiated to photoreceptors (Fig. 4D). The transplanted cells displayed robust Jaws-induced photocurrents, demonstrating the functionality of the microbial opsin in the host environment (Fig. 4E). The measured photocurrents peaked at 575 nm and showed fast kinetics (Tau_ON_ < 10ms) (Fig. 4, F-H), reflecting the response properties of Jaws. At the ganglion cell level, we observed ON- and OFF responses from different ganglion cell types, which shows that Jaws- driven signals from transplanted photoreceptors were transmitted via second order neurons (Fig. S6) to ON- and OFF-ganglion cells (Fig. 4I and Fig. S7). The light intensity requirements were below the safety threshold for the human retina ^18, 19^ (Fig. 4J and Fig. S7). After transplantation of control human donor cells, expressing GFP only, no light responses were detected (Fig. 4K), as expected from the absence of OS.

**Fig. 4.**
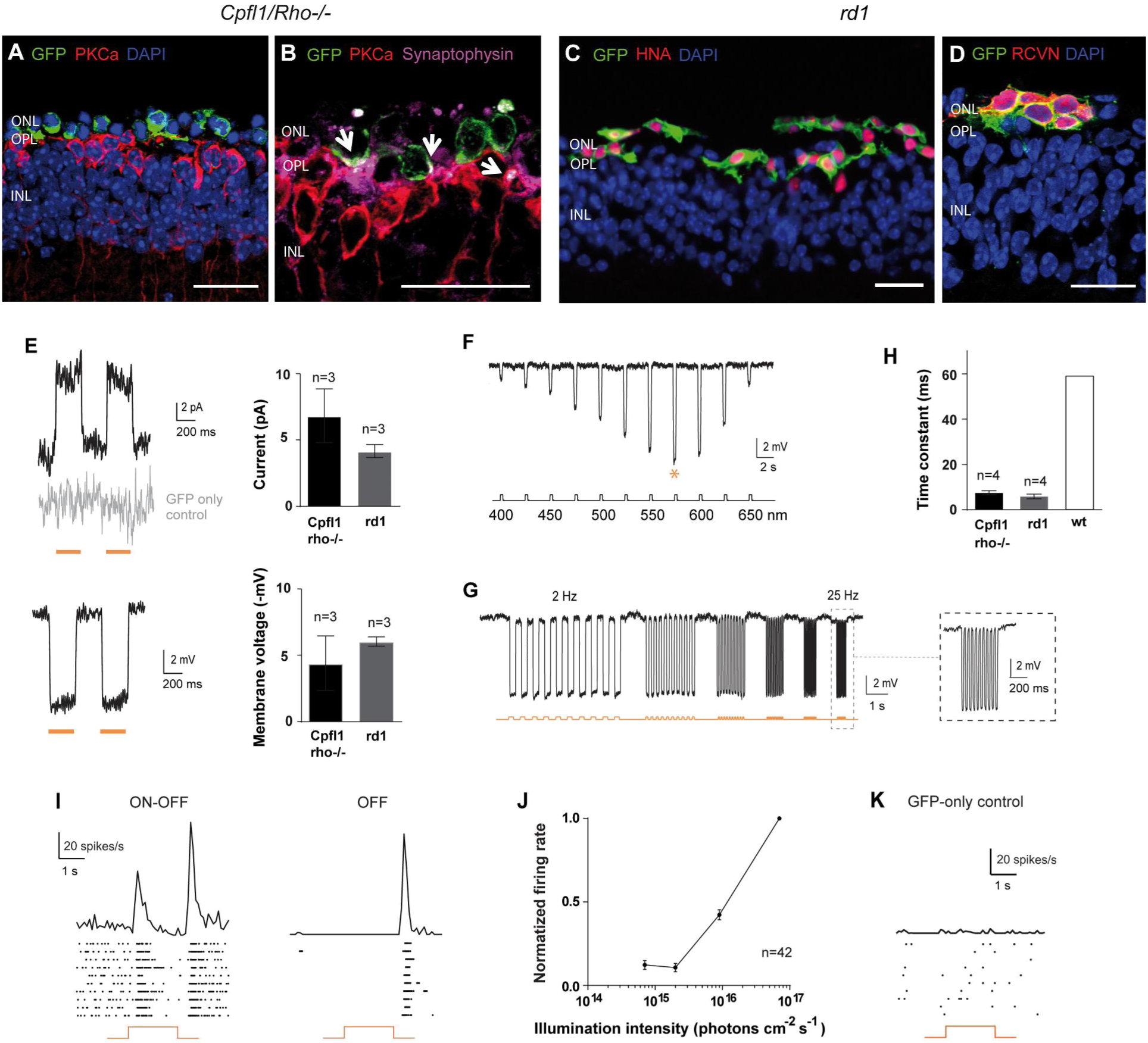
Transplanted photoreceptors, derived from hiPSCs, integrate into the retina of blind mice and display Jaws induced light responses that are transmitted to retinal ganglion cells. Immunofluorescence analysis on vertical sections 4 weeks after transplantation of Jaws-cone-treated *Cpfl1/Rho*^*-/-*^ (A,B) and *rd1* (C,D) retinas. **(A)** Transplanted cells (green) overlie host PKCα bipolar cells (red), DAPI counterstaining (blue). **(B)** Immunofluorescence against GFP (green), PKCα (bipolar cells, red) and synaptophysin (synapses, magenta). Arrows point to synaptic connections. **(C)** GFP+ Jaws-cones co-express Human Nuclear Antigen (HNA). **(D)** Transplanted GFP+ Jaws-cones also express RCVRN. Scale bars 20 µm. (E-I) Patch-clamp data from Jaws-cones after transplantation into blind mice, stimulation at 590 nm if not stated otherwise. **(E)** Left, representative photocurrents (top) and voltage hyperpolarization (bottom) after stimulation with 2 consecutive flashes, absence of the response in GFP-only cones shown in grey. Right, comparison of response amplitudes of Jaws-cones in different models (top, mean photocurrent peak; bottom, mean voltage peak). **(F)** Voltage action spectrum corresponding to a Jaws-expressing cell stimulated at wavelengths from 400 nm to 650 nm. Maximal responses were obtained at 575 nm. **(G)** Temporal properties: Jaws-induced hyperpolarization at increasing stimulation frequencies from 2 to 25 Hz. **(H)** Comparison of response rise time constant between Jaws-cones transplanted in *Cpfl1/Rho*^*-/-*^ and *rd1* models, and wt cones. (I-K) Averaged spike responses obtained from MEA recordings shown as PSTH and raster plots from a transplanted *Cpfl1/Rho*^*-/-*^ mouse. **(I)** Representative examples of two RGCs responding either with an ON/OFF or OFF-response (stimulation: 580 nm, 7 × 10^16^ photons cm^-2^ s^-1^). **(J)** Intensity curve. **(K)** Unresponsive cell from a control retina transplanted with GFP-only cones.

In conclusion, by using stem cells equipped with a microbial opsin, we went beyond the current limitation of an optogenetic approach which can only rescue the function of remaining ‘dormant’ cones ^24^. Here, we use the synergy of cell replacement and optogenetic therapy that allows to restore retinal *structure* with stem cells and visual *function* with microbial opsins. In a future perspective, optogenetically engineered hiPSC-derived cones could serve as donor cells for photoreceptor transplantation at late-stage retinal degeneration.

## Materials and methods

Description of methods and materials used are available as part of Supplementary information.

## Acknowledgments

We thank Thierry Leveillard for providing the *rd1* mice and Cheryl Craft for providing the hCAR antibody. We are thankful to Romain Caplette, Olivier Marre and Stéphane Deny for their help with the MEA recording and analysis. We thank Abhishek Sengupta for the construction of the light/dark box. We are grateful to Mélissa Desrosiers and Camille Robert for AAV productions and the fundraising department of the Vision Institute.

## Author contributions

MG optimized AAV mediated transduction of hiPSC-derived retinal organoids, performed culture, imaging, qPCR and histology, designed experiments and wrote the manuscript. ML performed in vivo injections, cell transplantation, MEA recordings, behavioral experiments, confocal microscopy, histology, designed experiments and wrote the manuscript. AC performed patch-clamp recordings and 2-photon imaging. LG generated hiPSC-derived retinal organoids, performed in vivo injections, cell transplantations, behavioral experiments, imaging, and histology. FR performed behavioral experiments. TF contributed to cell transplantation. GG and SR helped to optimize hiPSC cultures and differentiation protocols. SP and JAS provided scientific input, financial and administrative support. OG provided hiPSCs and gave feedback on the manuscript. MA provided *Cpfl1/Rho*^*-/-*^ mice, contributed to cell transplantation and gave feedback on the manuscript. DD and JD designed experiments and wrote the manuscript.

